# Biomechanics of venom delivery in South America’s first spitting scorpion

**DOI:** 10.1101/2024.07.25.605134

**Authors:** Léo Laborieux

## Abstract

Venom is a metabolically expensive secretion used sparingly in a variety of ecological contexts, most notably predation and defense. Accordingly, few animals use their toxins in ranged attacks, and venom-squirting behaviour is only known from select taxa. In scorpions, species belonging to two genera are known to spray venom when threatened, and previous work in *Parabuthus transvaalicus* shows that venom delivery depends on perceived levels of threat. Here, I describe *Tityus* (*Tityus*) *XXX* sp. nov., a new species of Buthid scorpion from Cundinamarca, Colombia. Remarkably, this species is capable of venom spraying, a first for both the genus and the South American continent. Using frame-by-frame video analysis and ballistic equations, I show that *T. XXX* sp. nov. employs not one, but two types of airborne attacks with dramatic differences in range and venom expenditure. Further, the new species uses an unusually large reserve of prevenom-like secretion for spraying, as opposed to the costly venom used by other spitting scorpions. In light of these key specializations, I propose that venom spraying convergently evolved in response to different selection pressures, laying the groundwork for future investigation.

## 1 Introduction

Venoms are exceptionally diverse biochemical mixtures employed in various ecological contexts. Venom has independently evolved over 100 times in a wide range of taxa (Morgenstern & King, 2013) and supports many complex behaviours such as predation, antipredator defense, mating (Sentenská et al., 2017), intraspecific communication (Post & Jeanne, 1983) and competition (Whittington & Belov, 2014), or habitat modification (Frederickson et al., 2005). Importantly, venom often serves multiple functions, and individual species may rely on venomous secretions in multiple ecological contexts. This versatility is enabled by specialized venom systems and, in many cases, associated with modulation of venom delivery. This modulation may be quantitative (“venom metering”, Hayes et al., 2002; Morgenstern & King, 2013), qualitative (i.e. achieved through differential toxin use), or both (reviewed in Schendel et al., 2019), and reflects selection pressures driving venom economy.

The outstanding diversity of venomous animals unequivocally demonstrates the ecological opportunity presented by venom, yet replenishing toxin reserves and maintaining a venomous apparatus also carries significant costs. Previous studies have shown increased metabolic rates in the days following venom delivery in pit vipers (McCue, 2006; Smith et al., 2014) and scorpions (Nisani et al., 2012), and hinted at prolonged metabolic costs of venom regeneration in both snakes (Oron & Bdolah, 1973) and araneomorph spiders (Boevé et al., 1995). Beyond this direct impact, indiscriminate use of venom reserves is also expected to negatively affect both defensive and predatory success by simply elevating the risk of depletion. Therefore, venom expenditure is thought to be “optimized” (Kuhn-Nentwig et al., 2011; Morgenstern & King, 2013): while remaining as conservative as possible, it is adjusted based on various factors that can be broadly summarized as (i) the condition of the venomous animal and its venom apparatus, (ii) the level of immediate threat posed by the target (predator or prey), and (iii) its specific susceptibility to the toxins.

Some venomous animals have adapted to use venom at range, avoiding direct physical confrontation with the target. While this potentially represents a considerable advantage, venom projection is an inherently inefficient strategy. It is thus expected that the energetic cost of spraying, rather than injecting, is offset by significant survivability benefits (Evans et al., 2019). This tradeoff remains poorly characterized, and the proportion of venomous animals able to project their secretions is small. Perhaps the best studied example of venom spraying comes from spitting cobras and rinkhals, in which venom projection independently evolved three times (Wüster et al., 2007; Panagides et al., 2017). Modified fangs enable these elapid snakes to spit venom up to a distance of 3m, targeting the eyes of potential antagonists (Bogert et al., 1943; Rasmussen et al., 1995; Westhoff et al., 2010). Upon contact, convergently up-regulated PLA_2_ toxins induce high cytotoxicity, intense pain and, in some cases, blindness by destroying the cornea (Kazandjian et al., 2021). This behaviour is enabled by a suite of adaptations, such as the direction and shape of the discharge orifice (Bogert et al., 1943; Wüster & Thorpe, 1992), and the presence of ridges in the venom channel reducing pressure requirements for expulsion (Avella et al., 2021; Triep et al., 2013).

Airborne delivery of venomous secretions is not exclusive to snakes, and scorpions belonging to two genera are known to spray venom defensively. While the “spitting” behaviour of *Hadrurus* species (family Hadruridae) unfortunately appears to have been overlooked (see Ythier & Stockmann, 2010), that of *Parabuthus* scorpions (family Buthidae) has received more attention. Similar to other scorpions, the venom of *Parabuthus transvaalicus* Purcell shifts in composition over successive stings (Inceoglu et al., 2003; Lira et al., 2017; Yahel-Niv & Zlotkin, 1979), transitioning from translucent, painful, K+-rich prevenom to a potent and peptide-rich secretion (Inceoglu et al., 2003). This has been hypothesized to represent passive venom modulation mediated by non-uniform storage of venom components in the venom gland (Schendel et al., 2019). Indeed, pre-venom invariably precedes the comparatively costly venom, which is thus reserved for high-threat situations (Nisani & Hayes, 2011). When their metasoma (“tail”) is grasped by an aggressor, individuals may spray venom out of their aculeus and engage in rapid movement of both the metasoma and telson. This thrashing widens the stream, likely increasing the odds of successful contact with sensitive tissues (Nisani & Hayes, 2015). The white venom, used as a toxungen, acts similarly to those of spitting cobras, causing intense pain and blindness (Newlands, 1974). But while the venom spraying behaviour of *Parabuthus* is now well characterized, considering these species as the only spitting scorpions has precluded comparative approaches that could shed light on the evolution of this unusual strategy.

Here, I describe the first South American scorpion capable of venom spraying and characterize its venom delivery. The recent discovery of a population of *Tityus (Tityus*) scorpions in the Cordillera Oriental region of Colombia, and its morphological comparison to known members of the genus, warrants the description of a new species, *Tityus (Tityus) XXX* sp. nov. Importantly, this scorpion’s ability to spray venom, a first for the genus, represents a prime opportunity to compare its venom delivery strategy to that of *Parabuthus* species. To do so, I videorecorded over 40 airborne attacks at high frame rate. Using frame-by-frame analysis and ballistic equations, I estimate the velocity, trajectory and range of the toxungen employed by the new species. In addition, using the same tracking data, I calculate approximate volume expenditure and make predictions about the volume of venom reserves.

## 2 Materials and methods

### Experimental subjects

The scorpions used for the behavioural study were 10 juveniles (prosoma length 3.5 - 6.8 mm) collected at the type locality. Unfortunately, no adults could be used as the male holotype was the only mature individual collected despite considerable sampling. Specimens were kept in large clear plastic cups (height x diameter: 13 × 10 cm) with humidified substrate and hides. These cups were placed near a window in an uninsulated room, ensuring the subjects were exposed to a photoperiod and temperature fluctuations identical to wild conditions. Food was offered every six days (see below).

### Venom-spraying experiment

Videos were recorded using an Apple iPhone 12 mini at 240 frames per second. The smartphone was chosen for (i) its high frame rate capability, (ii) its built-in flash providing constant and standardized illumination, (iii) the low ground clearance achieved when the device was used upside-down and (iv) quick repositioning between takes. Individual scorpions were gently transferred to the experimental box, a clear rectangular plastic container lined with a sheet of (5mm) grid paper. After five minutes of undisturbed acclimation, the camera was positioned to the side, perpendicular to the scorpion’s antero-posterior axis and level to the bottom of the enclosure. Shortly after recording started, scorpions were pinned to the bottom using a vertically oriented plastic straw of known diameter (5.6 mm) to trigger a defensive response. In practice, pressure was repeatedly applied to the carapace by briefly (∼0.5 – 1s) pushing the straw downward. Individuals that fled were allowed to settle down once more, and the process was repeated until stinging/squirting was abandoned entirely in favor of escape responses. The relatively soft straw was chosen to avoid harming the individuals, but also because of its cylindrical cross-section. This had the significant benefit of providing a reliable scale directly in the sagittal plane of the scorpion, reducing bias from perspective effects.

The trial group was tested every 6 days, and food was offered in the form of small insects (Orthoptera/Blattodea, size = approx. 2x prosoma length) immediately after each of 5 experimental sessions. Individuals that molted were not retried to let their exoskeletons harden safely. Specimens 8 and 9 were fixed to serve as paratypes before the 4^th^ and 5^th^ experimental sessions, respectively. The remaining specimens were released where they were collected after the 5^th^ session.

### Analyses

Frame-by-frame video analysis was performed in Tracker V6.1.6. Here, I define a venom “pulse” as any uninterrupted ejection of venom with no regard for its specific characteristics. Venom pulses that were immediately stopped by the straw or the enclosure’s wall were not analyzed, since this prevented spatial tracking over multiple frames, but were nonetheless recorded. Similarly, analysis was not attempted when pulses were out of focus or severely misaligned.

To extract the characteristics of venom pulses, videos were converted to 240 fps .mp4 files and opened in Tracker. A spatial scale was set a few frames before the first pulse based on the diameter of the straw used for stimulation (calibration stick tool). The scale was not reset between pulses, as the scorpions generally moved minimally until they decided to flee. The scale was, however, reset whenever the camera had to be repositioned. Tracking was performed using the point mass tool on the forwardmost visible portion of the projected venom on consecutive frames. For a given venom pulse, the initial projection angle was measured by setting the origin of the reference frame on the first tracked point and recording the angular position of the second. The orientation of the x axis was set depending on that of the scorpion (left or right), such that an exactly forward-projected venom pulse had an initial angle of 0° (vertically projected: 90°). For sustained (> 7 frames) pulses, the minimal and maximal angles of expulsion were also identified and recorded. Each of these measurements was made by placing the origin on the tip of the aculeus and recording the angular position of the venom on the next frame. The initial height of projection was determined by placing the origin of the reference frame in the plane of the scorpion’s sternites and recording the vertical coordinate (y) of the first tracked point. The width of the venom stream was measured at the tip of the aculeus using the tape measure tool. Here, similar to Nisani & Hayes (2015), I consider that negative and positive parallax errors (introduced whenever the venom pulse propagated outside of the sagittal plane of the scorpion) had equal incidence, likely self-compensating.

The trajectories and ranges of venom pulses were estimated using ballistic equations. Aerodynamic drag was not considered as air resistance was unlikely to significantly affect estimates: the cross-sectional area of the venom stream was small (approx. 0,05 *mm*^2^ assuming a circular cross-section) and the velocity of the projections was relatively low. Under these simplifying assumptions, the trajectory of a projection of velocity *ν* with initial angle *θ*_*i*_ and height *h*_*i*_is given by:

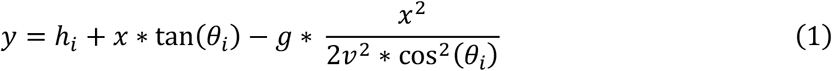

Where *g =* 9,81 *m*. s^-2^. The range of the secretion, or the distance it will travel before falling back to the ground, is given by:

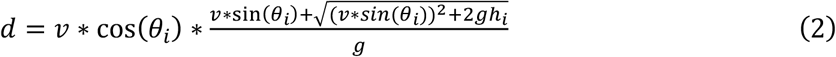

And the apex of the trajectory can be calculated as:

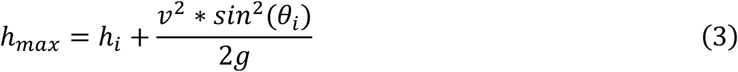

Tentative volume estimates were generated by modelling the projected secretion as a cylinder of constant diameter whose length depended on the velocity and duration of the venom pulse. The measured width of the venom stream exiting the aculeus was stable across individuals and venom pulses, with an average value of approx. 250 µm (Table 1). The projected volume was thus modelled as:

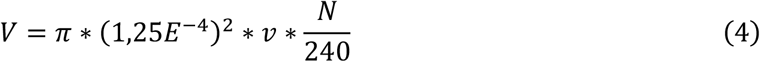

**Table 1.** Venom stream width measurements for venom flicks and venom sprays. N/As indicate missing data for specific (venom pulse type/specimen) combinations.

| Specimen | Stream width (flick) | Stream width (spray) |
| --- | --- | --- |
| 1 | 254 | 272 |
| 2 | N/A | 243 |
| 3 | N/A | 231 |
| 4 | 253 | N/A |
| 5 | 250 | 243 |
| 6 | 278 | N/A |
| 7 | 236 | 231 |
| 8 | N/A | N/A |
| 9 | 247 | 239 |
| 10 | 281 | 241 |
| Mean | 257 | 242.85 |
| Overall mean | 249.92 |  |

Where *ν*is the velocity (*m*. s^-1^) estimated for the venom pulse and *N*is its duration in frames.

Statistical analyses were conducted in R (R Core Team, 2023) and plots were produced using the *ggplot2* package (Wickham, 2011). To analyze differences between venom pulse types, linear mixed-effects models (LMM) were fitted using the *lmer()* function of the lme4 package (Bates et al., 2015). Venom pulse type was the only fixed effect, and models were built to investigate its effect on ejection velocities, ranges, vertical apexes and ejected volumes. To also test whether distinct venom pulse types were initiated at different heights while controlling for individual size, I calculated a scaled metric equal to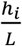, where L is the carapace length for the individual considered. This parameter was analyzed using a linear model with an identical fixed effect structure. Because individuals were tested multiple times, and to avoid pseudoreplication, individual ID was included as a random factor. In the case of the velocities, ranges and apexes, however, including this random factor resulted in a singular fit, indicating that inter-individual variation was negligible. In these instances, linear models were refitted using the *lm()* function, thus omitting the specimen-level random factor. The significance of the fixed effect was assessed using the *Anova* command of the car package (Fox & Weisberg, 2019) and contrast assessments were generated using the *emmeans* function of the emmeans package (Lenth et al., 2023). To investigate the relationship between the type of venom pulse and the initial angle of projection while controlling for pseudoreplication, angular distributions were compared using a mixed-effects model with a von Mises distribution as implemented in the *brm* function of the brms package (Bürkner, 2017), with a normal prior (µ = 0, σ^2^ = 30°). Convergence of parameter estimates was assessed based on R-hat values. Bayes factors (BF) were computed as implemented in the *bayesfactors_parameters* function of the bayestestR package (Makowski et al., 2019).

### Systematics

Photographs were taken using a Fujfilm X-Pro 3 digital camera paired with a Laowa 65mm f/2.8 2X APO macro lens. Photographs were taken as long exposures, and light sources (white or UV) were rotated around the subjects during exposure to minimize shadows. Figures constructed from several photographs were produced in Adobe Photoshop. Measurements follow Stahnke (1970) and were made using a vernier caliper. The lengths of individual metasomal segments are given as their full anatomical length as revealed when the metasoma is fully flexed. Anatomical terminology mostly follows Vachon (1952) and Hjelle (1990). Trichobothrial notations follow Vachon (1974). Terminology specific to the pectinal basal piece and walking leg telotarsi is outlined in Moreno-González et al. (2021). The holotype and paratypes studied herein are deposited at the Museo de Historia Natural C.J. Marinkelle (Universidad de los Andes) in Bogotá, Colombia.

## 3 Results

### 3.1 Venom-spraying experiment

A total of 46 venom pulses were videorecorded at high frame rate, of which 38 could be analyzed frame by frame. The scorpions invariably exhibited defensive behaviour when compressed by the straw. All but one (90%) of the individuals in the trial group projected venom at least once over the span of the experimental period. Two distinct types of venom pulses could be characterized: (i) venom flicks, defined here as short projections consisting of a single droplet and (ii) venom sprays, sustained projections of venom with distinct biomechanical characteristics. In total, 14 flicks and 24 sprays were analyzed.

Venom flicks and venom sprays were easily discriminated based on important differences in duration and velocity **(Fig. 2A)**. During flicks, venom expulsion was succinct (≤ 1 frame / 4.16 ms, hereafter considered to equal 1 frame) while it was considerably longer during sprays, albeit with high variability (Mean ± SD: 28.2 ± 22.4 ms). Variation in spray duration spanned a whole order of magnitude (min - max: 8.3 - 95.8 ms). Duration was not the only discriminatory metric between the two pulse types, however, as expulsion velocity was significantly higher for venom sprays (1.32 ± 0.35 m.s^-1^) than for venom flicks (0.41 ± 0.22 m.s^- 1^) (F = 77.448, df = 1, p = 1.681E-10; “emmeans” contrast assessment: ß±SE = -0.915 ± 0.104; **Fig. 2A**). The highest expulsion velocity recorded was 1.89 m.s^-1^ (specimen 5). The velocity ranges for both pulse types did not overlap, except for one abnormally fast flick (specimen 9; *ν* = 1.02 m.s^-1^). The latter was not classified as a spray based on its duration.

**Fig. 1.**
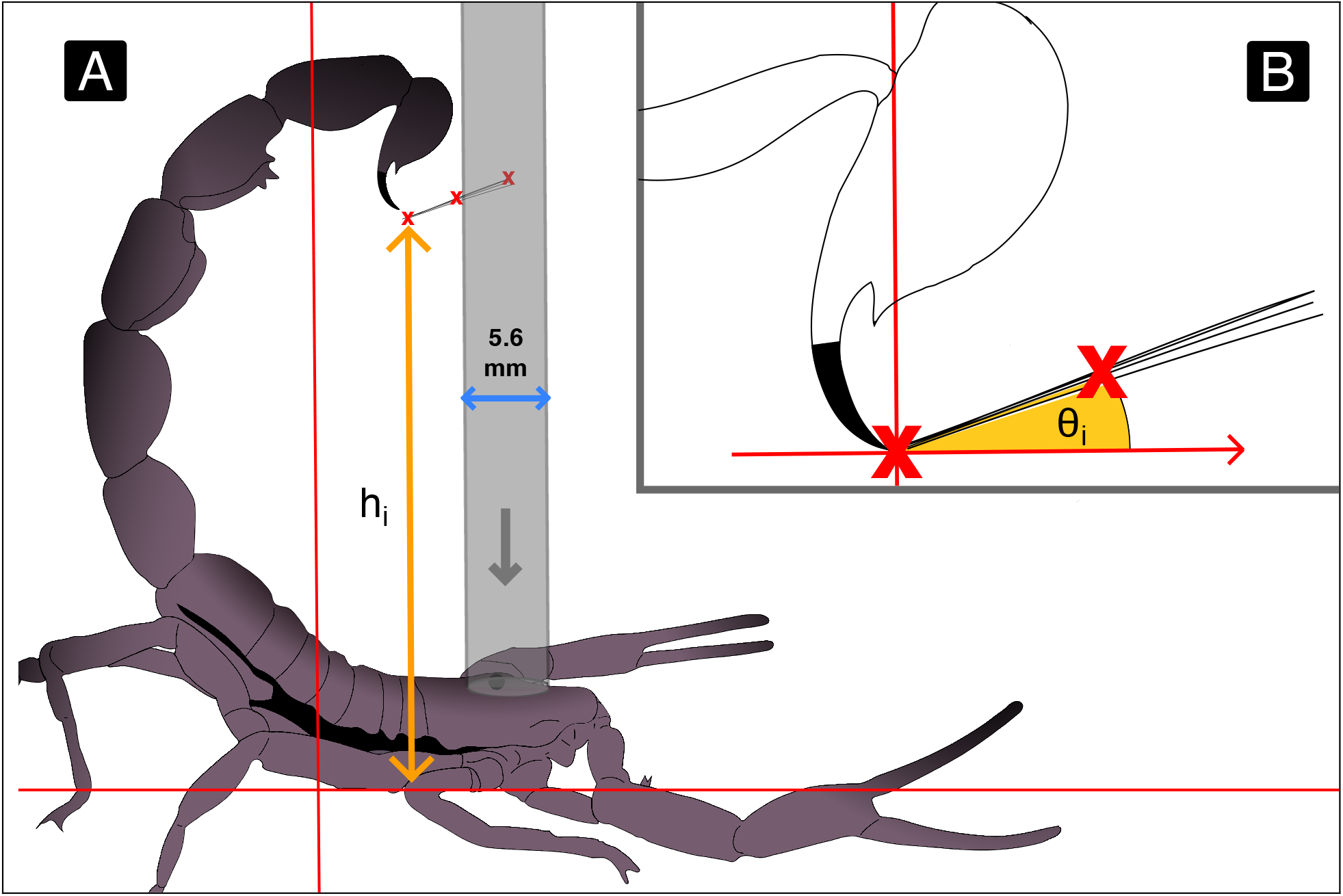
Summary of video data extraction for venom pulses. **A**. The spatial scale (blue arrow) is set perpendicular to the straw (5.6 mm diameter). The initial height *h*_*i*_ (orange arrow) is the difference in height between the sternites of the scorpion and the first tracked point. Red lines indicate the position of the reference frame. **B**. The initial angle *θ*_*i*_(orange arc) is recorded as the angular position of the second tracked point relative to the first. Red crosses mark tracked positions on consecutive frames.

**Fig. 2.**
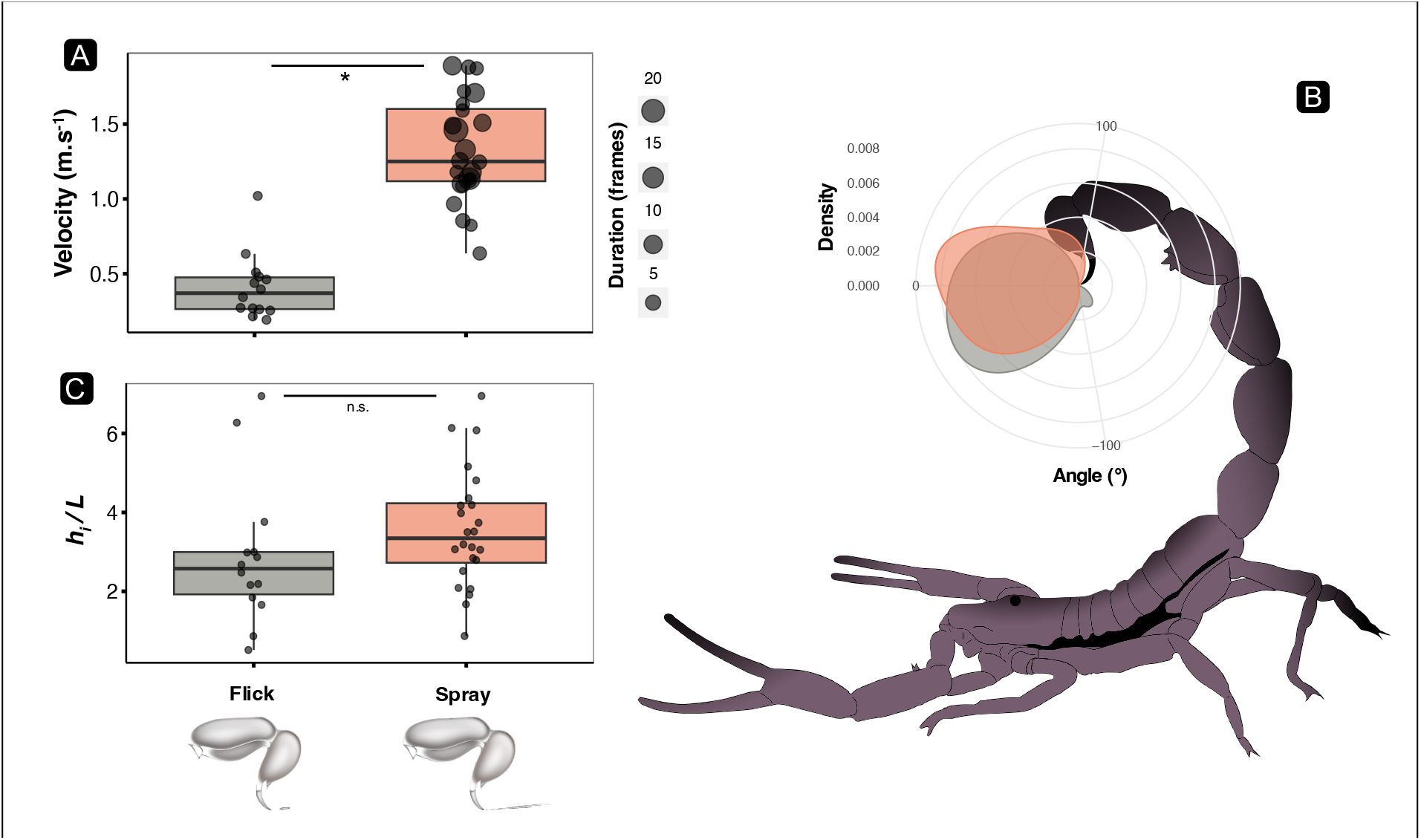
Characteristics of venom flicks (gray, N = 14) and sprays (orange, N = 24) as extracted from high-speed video. A: Observed projection velocities for venom flicks and venom sprays. Point size indicates pulse duration in frames. Together, these characteristics served as the basis for discriminating flicks and sprays. B: distributions of the initial angles of projection for flicks and sprays. C: Adjusted initial projection heights 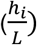. A star (*) denotes statistical significance.

Venom pulses were generally directed forward **(Fig. 2B)**, irrespective of type. One venom flick (specimen 5) was distinctly backwards-directed (*θ*_*i*_ = -123.7°) due to metasomal movements, while two venom sprays were directed vertically (specimen 1: *θ*_*i*_ = 90.0°; specimen 10: *θ*_*i*_ = 90.3°). There was little evidence in favor of initial projection angles (**Fig. 2B)** differing significantly between the two pulse types (Posterior mean effect of pulse type: 0.19 [95% CrI: - 0.07 to 0.45]; BF = 1.29), though the angular distributions showed a positive (*θ*_*i*_> 0°) mode for sprays and a negative mode (*θ*_*i*_< 0°) for flicks. Long sprays (> 7 frames / 29.1 ms) showed considerable deviation from their initial direction due to metasomal movements. For these sprays, the angular range of projection was narrow on average, but variable (49.9 ± 29.0°, N=9), with a maximum of 97.0°. Overall, these movements appeared similar to those associated with regular stings, though this cannot be affirmed without precise spatial tracking (see Coelho et al. (2017) for an overview of scorpion stinging motion). Further, there was partial support for differences between the adjusted initial projection heights 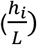 of venom flicks and sprays. While the data distributions suggest that venom sprays were initiated higher than venom flicks on average (**Fig. 2C)**, this was not statistically supported (X^2^ = 1.7496, df = 1, p = 0.1859; “emmeans” contrast assessment: ß±SE = -0.427 ± 0.33).

Together, *ν, h*_*i*_ and *θ*_*i*_were used to generate the estimated trajectories shown on **Fig. 3** (Eq. 1). The range of venom flicks (Mean ± SD: *d* = 1.54 ± 1.56 cm) was significantly lower than that of sprays (9.53 ± 10.88 cm) (F = 7.3631, df = 1, p = 0.01015; “emmeans” contrast assessment: ß±SE = -0.0799 ± 0.0294) and the maximum ranges recorded for flicks and sprays were 4.93 cm (specimen 7) and 36.10 cm (specimen 5) respectively. Similarly, the vertical reach *h*_*max*_ was significantly higher for sprays than for flicks (X^2^ = 6.9325; df = 1, p = 0.008464; “emmeans” contrast assessment: ß±SE = -0.0341 ± 0.0134). The highest vertical reach was achieved by a nearly vertical venom spray (specimen 10, *θ*_*i*_ = 90.3°, not plotted), with *h*_*max*_ = 20.12cm. The highest forward projected venom spray (specimen 5) reached 15.95 cm above ground. Consistent with other parameters, there was significant variability in *h*_*max*_ for both flicks (Mean ± SD: *h*_*max*_ = 2.35 ± 1.24 cm) and sprays (5.76 ± 4.73 cm).

**Fig. 3.**
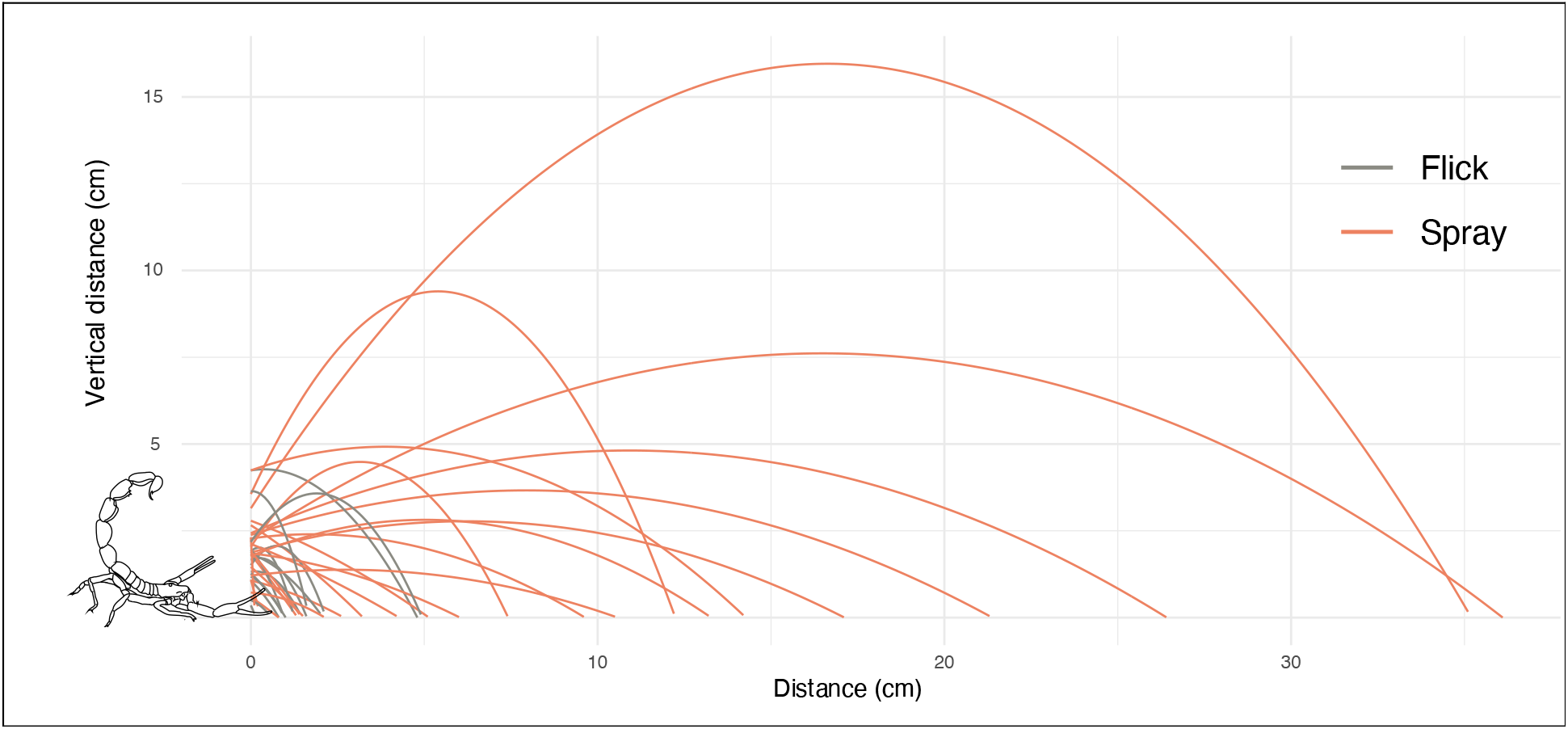
Venom pulse trajectories for flicks (gray) and sprays (orange) as calculated from equation (1) (two vertical venom pulses not represented for readability). Adult scorpion to scale.

In accordance with the biomechanical characteristics of both venom pulse types, the volume estimates for venom flicks and venom sprays showed dramatic differences (**Fig. 4)**. The estimated volume of venom expelled during venom sprays was significantly larger than that associated with venom flicks (X^2^ = 17.793; df = 1, p = 2.463E-05; “emmeans” contrast assessment: ß±SE = 1.81 ± 0.442). The mean volumes of venom expelled during venom sprays and flicks were 1.98 µL (SD: 1.60 µL) and 0.084 µL (SD: 0.045 µL) respectively. There was a 175-fold difference in estimated volume between the smallest venom flick (0.039 µL) and the largest venom spray (6.88 µL).

**Fig. 4.**
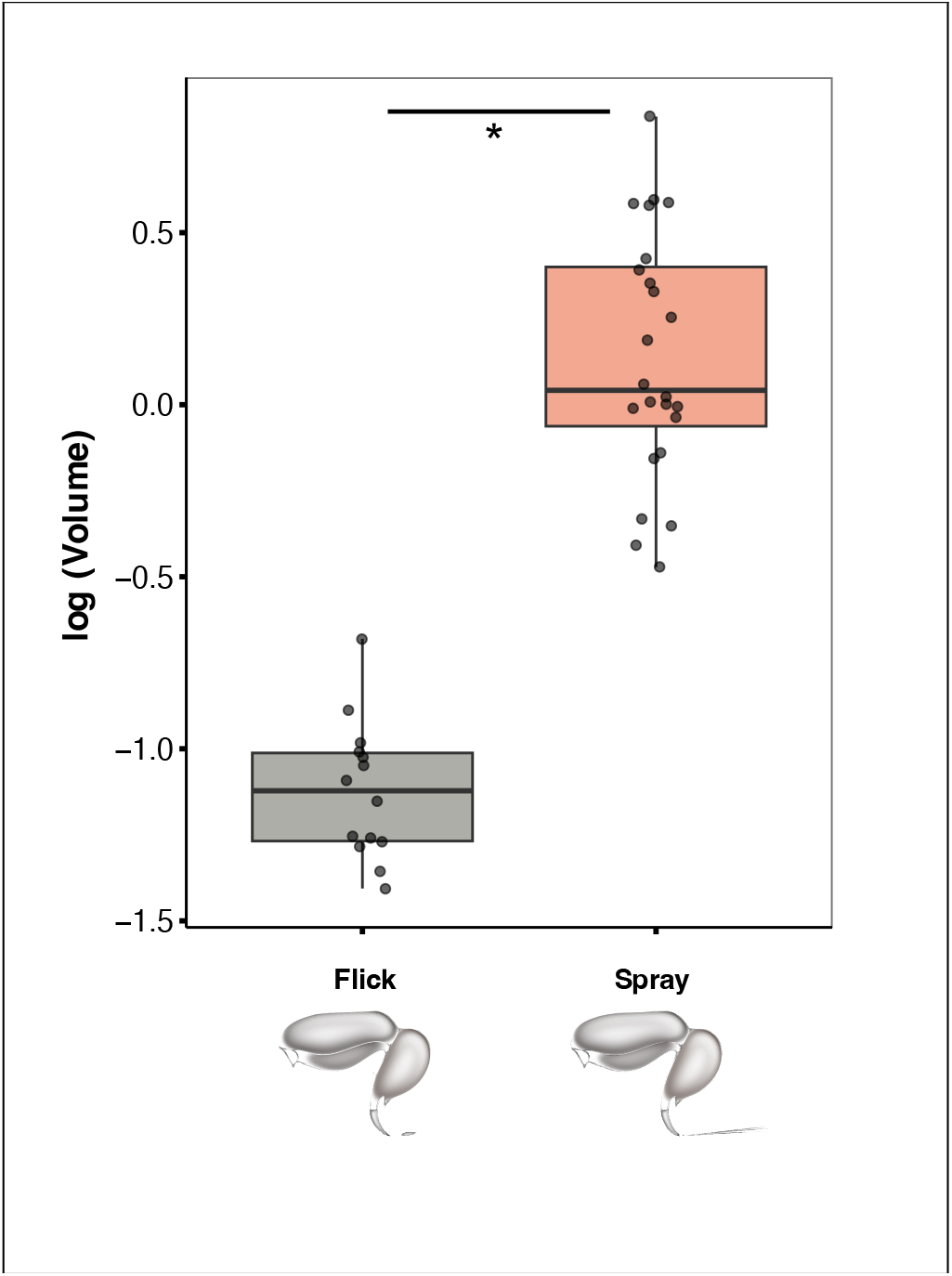
Expelled volume estimates for venom flicks (gray) and venom sprays (orange). Data plotted as log_10_-transformed to facilitate visualization. Star indicates statistical significance (test conducted on non-transformed data).

### Secretions used for venom pulses

When provoked, individuals were observed to employ two distinct types of secretion for defense. The first, a clear secretion **(Fig. 5A)**, was used extensively for defensive stings, venom flicks and venom sprays. This prevenom-like secretion was used for as many as five successive venom pulses (specimen 5, prosoma length = 6.1 mm), totaling an estimated expelled volume of 11.08 µL. Additionally, a second, more opaque secretion was observed on two occasions. Interestingly, this was in the case of both a venom spray (**Fig. 5B**, untested specimen) and a defensive sting (individual 7, prosoma length = 5.8 mm). In the case of individual 7, white venom was expelled after 6 venom pulses, though one (a short, 2-frame venom spray) could not be accurately tracked. The 5 tracked (transparent) venom pulses added up to an estimated volume of 9.45 µL, consistent with a large, approx. 10µL reserve of clear secretion. Collection of venom samples was attempted at a later stage, but no opaque secretion could be further obtained from the trial group.

**Fig. 5.**
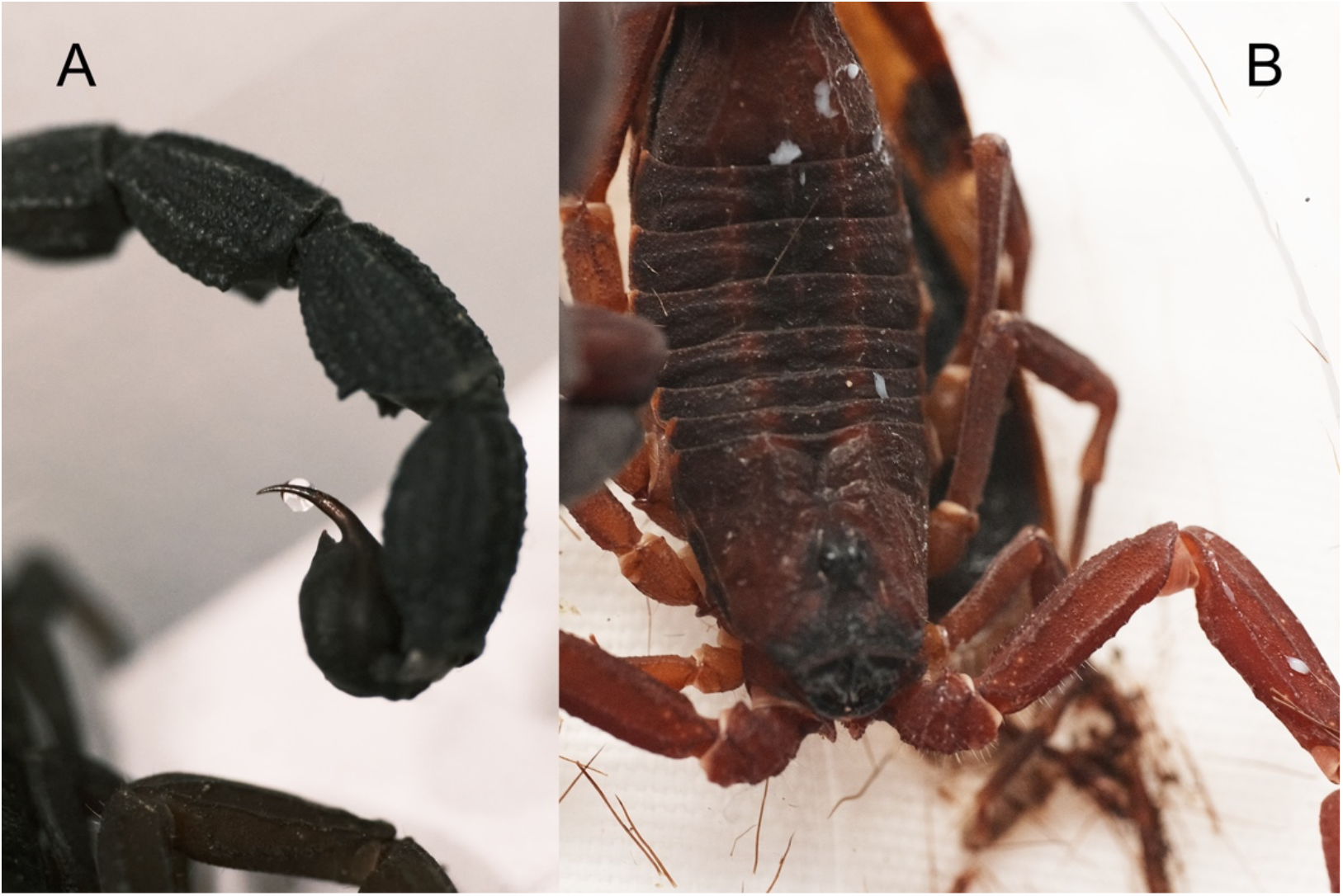
Distinct secretion types employed for venom pulses. A: clear, prevenom-like secretion. B: venom (white)

### 3.2 Systematics

Family **Buthidae** C. L. Koch, 1837

Genus ***Tityus*** C. L. Koch, 1836

Subgenus ***Tityus***

***Tityus* (*Tityus*) *XXX* sp. nov**.

Holotype, adult ♂, Colombia, Cundinamarca, La Vega, Estación Experimental JCM (4.955239,-74.379291), 1350m a.s.l., December 20, 2023, Léo Laborieux, deposited at the Museo de Historia Natural C.J. Marinkelle (Universidad de los Andes). Catalog number : ANDES-IN-8788

Paratypes, 1 subadult ♀ + 1 subadult ♂, Colombia, Cundinamarca, La Vega, Estación Experimental JCM (4.955239, -74.379291), 1350m a.s.l., December 20, 2023, Léo Laborieux, deposited at the Museo de Historia Natural C.J. Marinkelle (Universidad de los Andes). Catalog numbers : ANDES-IN-8789 (♂), : ANDES-IN-8790 (♀)

#### Etymology

The specific epithet refers to Greek mythology’s XXX, for his prowess as a spear wielder.

### Diagnosis

Scorpion of moderately large size for the subgenus, with a total size of 66,4 mm for the adult male holotype. General coloration dark red; carapace with median ocular tubercle and anterior margin black. Telotarsi of leg pairs I-IV with 2 discrete ventro-submedian rows of setae (type II). Axial carina on mesosomal segments I-VII subtle with feeble granulation. Paraxial carinae vestigial in mesosoma. Intercarinal spaces moderately granular on I-VI, and strongly granulated on VII. Pectines with 16-18 teeth in male holotype, 15-16 in female paratype. Basal middle lamellae dilated. Basal pectinal piece without a glandular region. All metasomal carinae feebly raised and subcrenulate. Metasomal segment II with 2 vestigial lateral carinae occupying its distal fifth. Metasomal segment IV with 1 prominent spinoid posterior granule on dorsal carina, and 1-2 smaller ones on dorsolateral carina. Aculeus curvature low. Chela fingers black with yellow tips. Inner face of movable finger with 15 oblique rows of denticles. Secondary sexual dimorphism in pedipalps moderate with femur, patella and chela length/width ratios 3.42 - 3.90, 3.41 - 3.83 and 6.39 - 6.6 respectively in males 3.75, 3.68 and 5.13 respectively in female paratype. This species may also be distinguishable by particularly low UV-induced fluorescence.

### Description

Based on adult male holotype and subadult female paratype.

#### Coloration

Overall dark and reddish. Prosoma: anterior margin of carapace and carinae around the ocular median tubercle black. Intercarinal spaces dark with a red sheen. Mesosoma: darker than carapace, with two lighter paraxial lines. Metasomal segments progressively darkening from I to V, from the same dark reddish hue to black. Dorsal side of metasoma segments lighter than ventral sides. Telson lighter than V. Venter and pectines yellowish, lighter than the rest of the body. Pedipalps: uniformly reddish with the exception of black fingers. Tips of fixed and movable fingers yellow. Chelicerae black with deep red teeth. Little ontogenic change in coloration. Interestingly, this species shows only a low level of UV-induced fluorescence (**Fig. 11**), making night searches challenging. It is unclear whether this is unique to *T. XXX* sp. nov., or instead common to the whole subgenus.

**Fig. 6.**
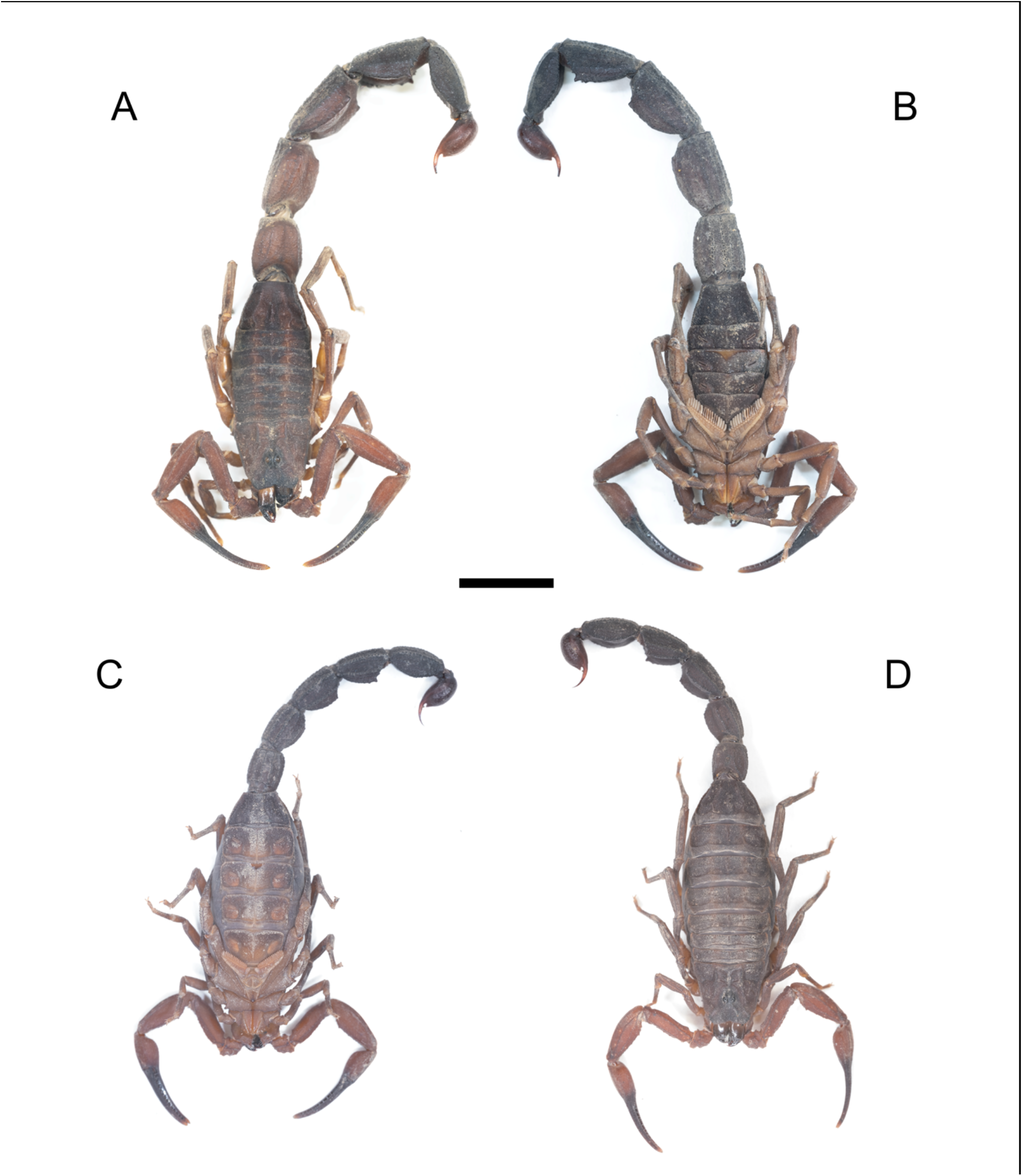
*Tityus (Tityus) XXX* sp. nov., habitus (Scale bar = 1 cm). **A-B :** adult ♂ holotype. **A**. Dorsal. **B**. Ventral. **C-D :** subadult ♀ paratype. **C**. Ventral. **D**. Dorsal.

**Fig. 7.**
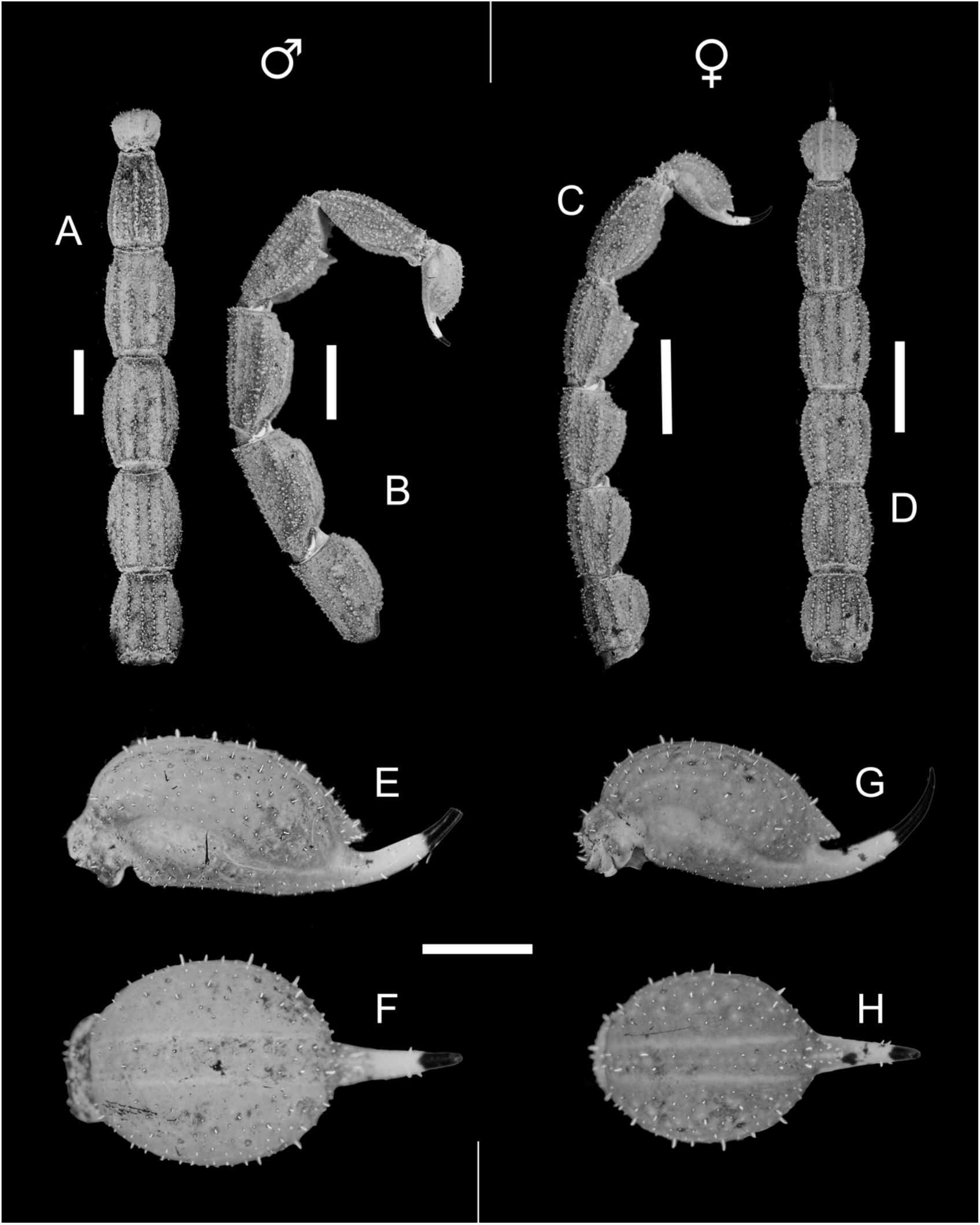
*Tityus (Tityus) XXX* sp. nov., metasoma. **A-B** : metasoma, ventral and lateral aspect, ♂ holotype. **C-D** : metasoma, ventral and lateral aspect, ♀ paratype. **E-F** : telson, lateral and ventral aspect, ♂ holotype. **G-H** : telson, lateral and ventral aspect, ♀ paratype. Scale bars = 5 mm (A-D) & 2 mm (E-H).

**Fig. 8.**
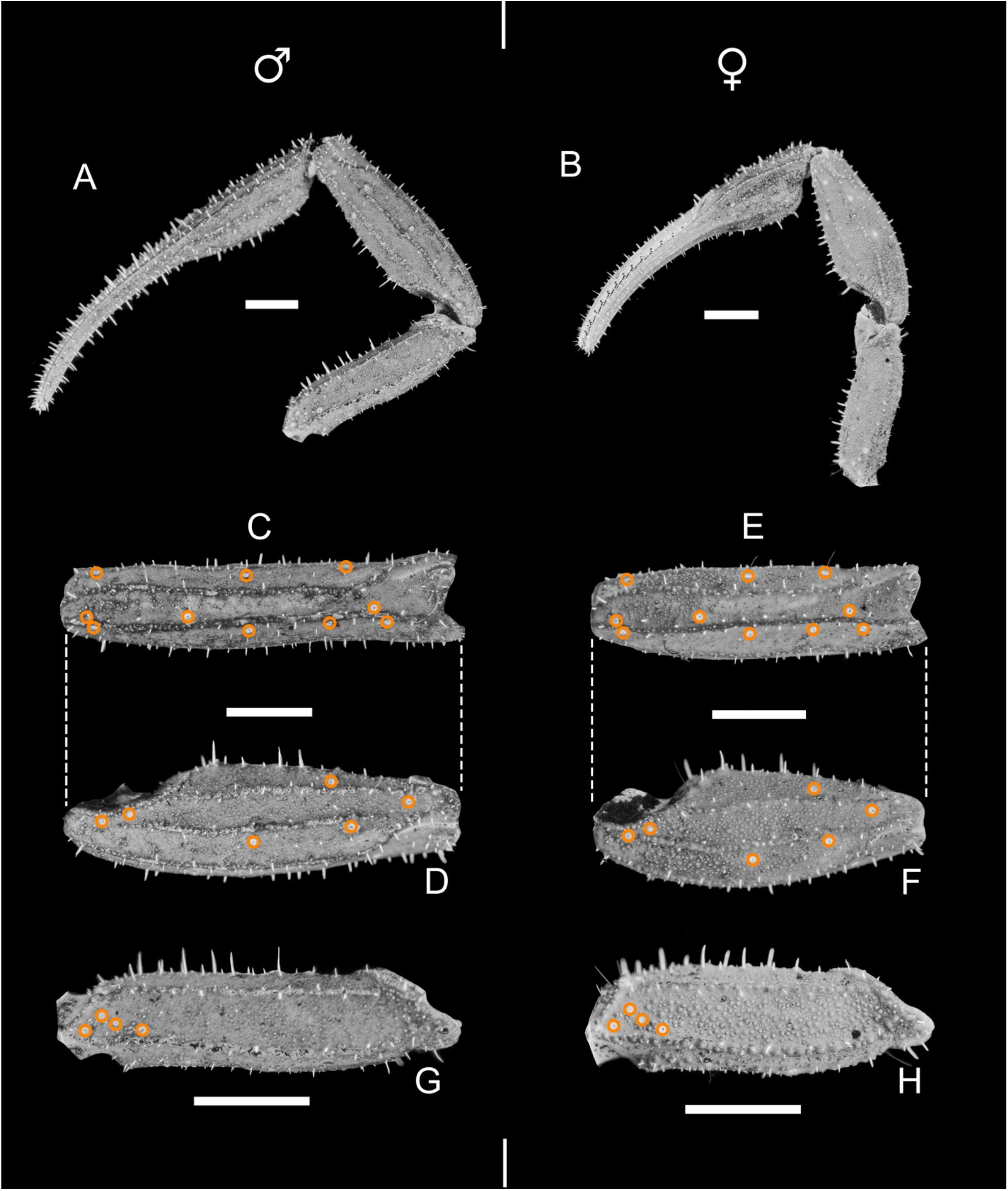
*Tityus (Tityus) XXX* sp. nov., pedipalps. **A**. ♂ holotype. **B**. ♀ paratype. **C-D** : patella, lateral and dorsal aspect, ♂ holotype. **E-F** : patella, lateral and dorsal aspect, ♀ paratype. **G-H** : femur, dorsal aspect. **G**. ♂ holotype. **H**. ♀ paratype. Orange circles mark the insertion points of the trichobothria. Scale bars = 2 mm

**Fig. 9.**
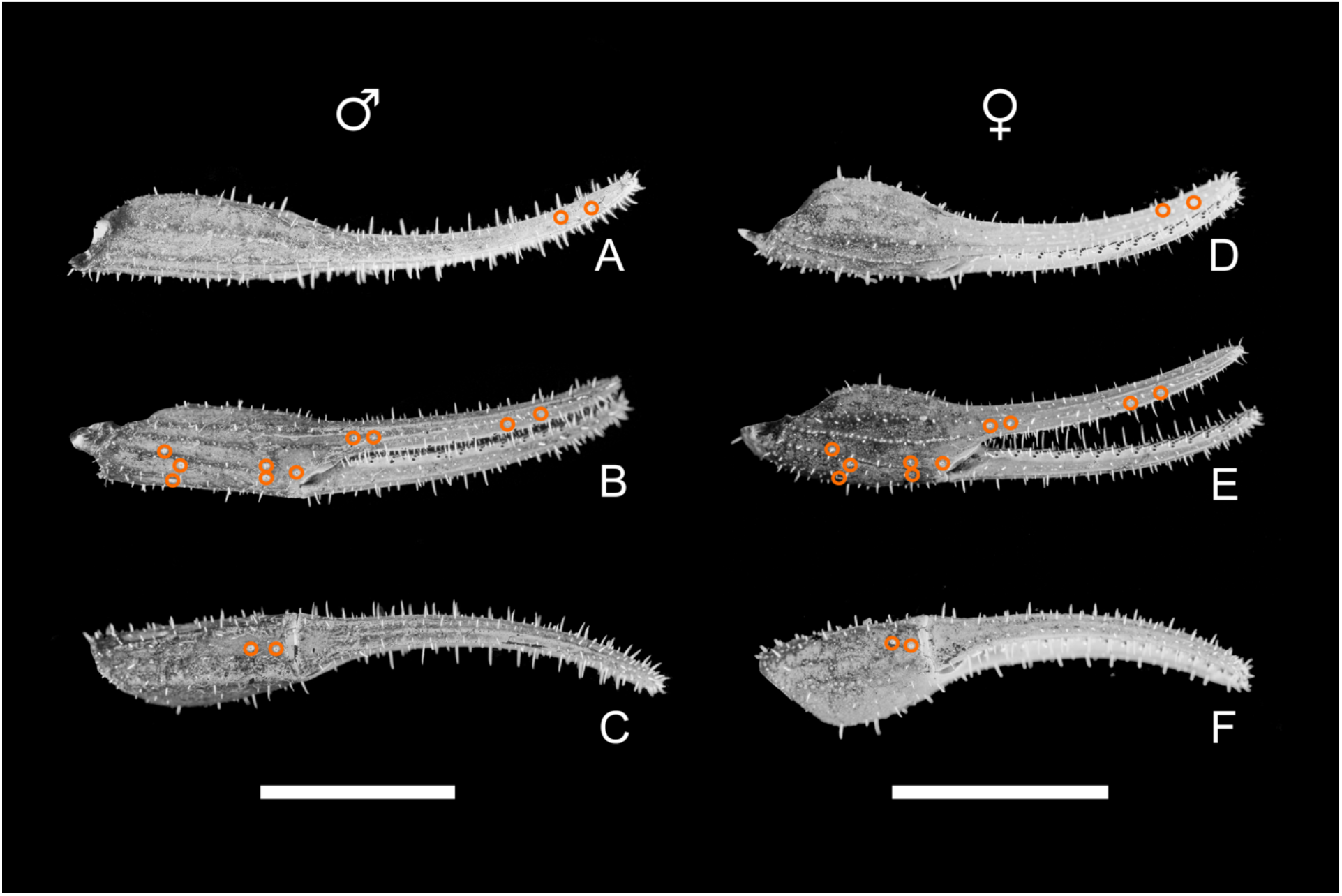
*Tityus (Tityus) XXX* sp. nov., chelae. **A-C :** dorsal, lateral, ventral aspects, ♂ holotype. **D-F :** dorsal, lateral, ventral aspects, ♀ paratype. Orange circles mark the insertion points of the trichobothria. Scale bar = 5 mm

**Fig. 10.**
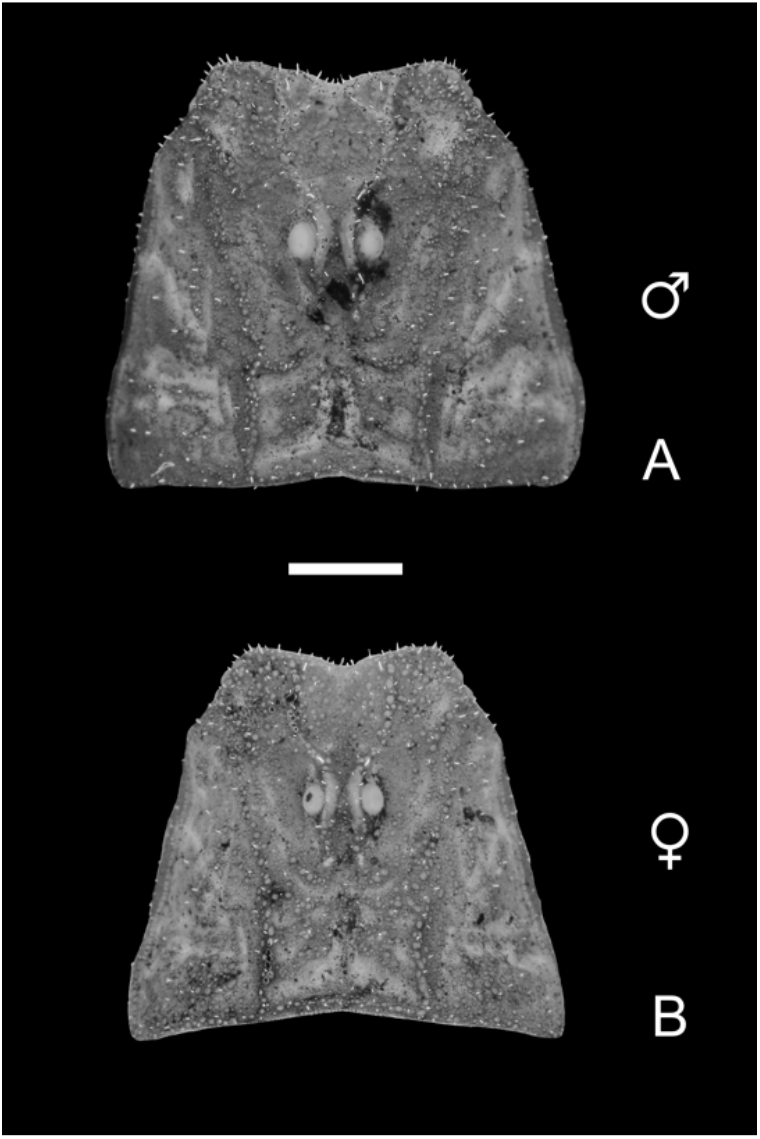
*Tityus (Tityus) XXX* sp. nov., carapace. **A :** ♂ holotype. **B :** ♀ paratype. Scale bar = 2 mm

**Fig. 11.**
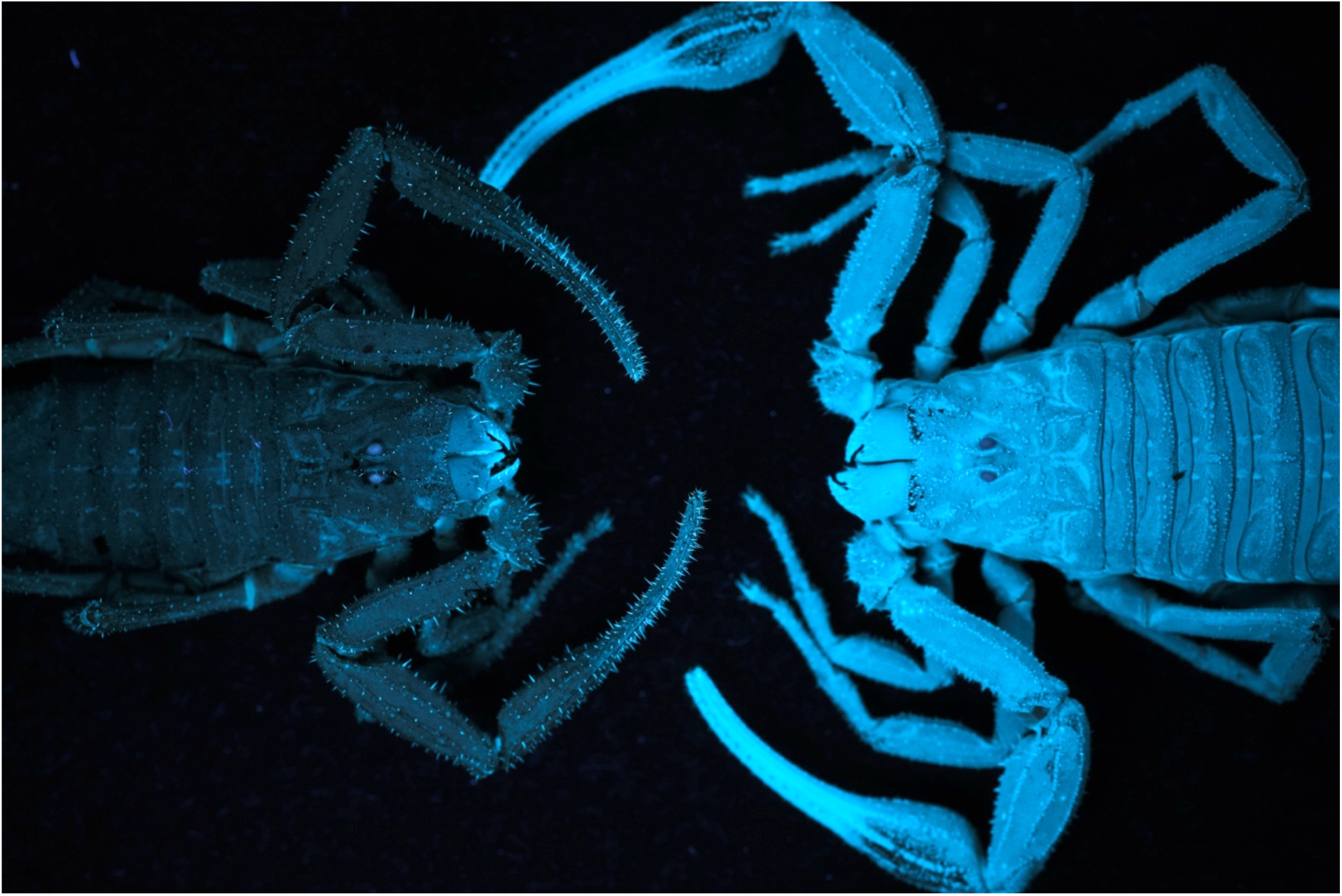
Comparison of UV fluorescence between the male holotype of *T. XXX* sp. nov. (left) and the female holotype of *T. (Atreus) icari* Laborieux, 2024 (right). The specimens were photographed under homogeneous UV lighting, and had received equivalent total UV exposure since collection.

#### Morphology

Carapace moderately to strongly granular, with the anterior margin presenting a marked median concavity. Furrows deep. Median ocular tubercle in a distinctly anterior position, with eyes separated by 1 ocular diameter and strong crenulate median ocular carinae becoming subcrenulate over the eyes. 3 pairs of lateral eyes, half the diameter of the median eyes. Sternum narrow, subtriangular. Telotarsi of leg pairs I-IV with 2 discrete ventro-submedian rows of setae (type II). Mesosoma: 1 central longitudinal carina in all tergites with feeble granulation; in I-VI represented by 1 row of granules in the posterior quarter and 2 rows of granules more anteriorly. 2 vestigial paraxial carinae on tergites I-IV. Tergite VII pentacarinate, with central longitudinal carina in anterior half. Lateral and paramedian carinae in VII joined proximally by a series of vertically organized and raised granules. Transversal carinae strong and distinct on tergites II-V, partially fused to posterior margin on tergite I. Intercarinal spaces moderately granular on I-VI, and strongly granular on VII. Venter with genital operculum split longitudinally; valves subtriangular.

Pectines: pectinal tooth count 16-18 in male holotype, 15-16 in female paratype. Basal middle lamellae dilated. Basal pectinal piece without a glandular region (based on female paratype). Sternites I-III feebly granular; IV and V moderately and strongly granular, respectively. Flattened triangle on the posterior margin of III, remarkably smooth. Sternite V with 2 + 2 crenulate longitudinal keels, i.e. (i) 2 paramedian keels on distal two thirds and (ii) 2 distinctly curved lateral keels occupying the second anterior quarter only. Spiracles oblique and of moderate length. Metasoma: all carinae feebly raised, subcrenulate. Setation weak. Segment I decacarinate; segment II with 8 carinae and 2 vestigial lateral carinae occupying its distal fifth. Segments III and IV with eight carinae. Segment V with seven carinae: 2 dorsal, 2 ventral, 1 median ventral and 2 paramedian ventral. Posterior granules of segment III weak, tubercular. Segment IV with dorsal and dorsolateral carinae displaying 1 strong and 2 - 3 weaker spinoid posterior granules, respectively. This spinoid granule on IV’s dorsal carina is prominent, representing its most obvious feature. Intercarinal spaces granular in all segments. Median dorsal depression present and moderate in V. Telson feebly granular, smooth dorsally. Aculeus short with low curvature. Subaculear tooth moderate, weakly spinoid. Cheliceral dentition characteristic of family Buthidae; median tooth of fixed finger strong and rather distinct. Trichobothrial pattern type A; orthobothriotaxic as defined by Vachon (1974). Dorsal trichobothria of femur arranged in α (alpha) configuration. Trichobothria *d2* present on the internal face of the femur (emigrated from dorsal face). Pedipalps with moderate overall setation, but fingers with strong setation. Femur setation concentrated on ventral side; patella with some short macrosetae scattered all over. Ventral side of manus almost devoid of setation; external and dorsal sides with a higher density of macrosetae than in patella. Fingers hairy, especially on internal sides with long macrosetae. External sides of fingers with increased setation distally. Femur with five crenulate and raised carinae, as well as an additional keel-like row of granules along the proximal third of its ventral side. Patella with 7 raised keels, all subcrenulate to smooth except for the 2 internal carinae, composed of spinoid granules. Intercarinal spaces moderately and feebly granular on femur and patella, respectively. Movable fingers with 15-15 oblique rows of denticles and strong, spinoid accessory and internal granules. First 2 proximal rows of denticles almost fused, but distinguishable based on their accessory and internal granules.

#### Comparisons

*T. XXX* shows affinities with the Colombian *T. (Tityus) lourençoi* Florez and *T. (Tityus) charalaensis* Mello-Leitão, with a generally dark-reddish coloration, male pedipalps slenderer than those of females, and fully parallel ventrolateral carinae on metasomal segments II-IV. It can however be easily distinguished from these by several key characters:

### -T. lourençoi

i. smaller overall size with the male holotype measuring 44.1 mm (66,4 mm in *T. XXX* sp. nov.)
ii. pronounced longitudinal keel on tergites (only slight in *T. XXX* sp. nov.)
iii. lower pectinal tooth count with 16-15 in male holotype (16-18 in *T. XXX* sp. nov.)
iv. 17 oblique rows of denticles on movable chela finger (15 in *T. XXX* sp. nov.)
v. longer pedipalps in male (pedipalp length/total length ratio 0,56 in *T. lourençoi* male holotype, 0,48 in *T. XXX* sp. nov.)
vi. broader patella in both sexes with length/width ratio 3.38 in male holotype and 2.86 in female paratype, compared to 3.41-3.83 (male) and 3.68 (female) in *T. XXX* sp. nov.
vii. broader chela in both sexes with length/width ratio 5.27 in male holotype and 3.87 in female paratype, compared to 6.39-6.6 (male) and 5.13 (female) in *T. XXX* sp. nov.
viii. shorter metasoma in both sexes with metasoma length/total length ratio 0,57 in male and 0,52 in female (0,66 and 0,58 in *T. XXX* sp. nov., respectively)
ix. spinoid posterior granules on segments I-IV (only on IV in *T. XXX* sp. nov.)
x. other morphometric values, see **table 2**

**Table 2.**
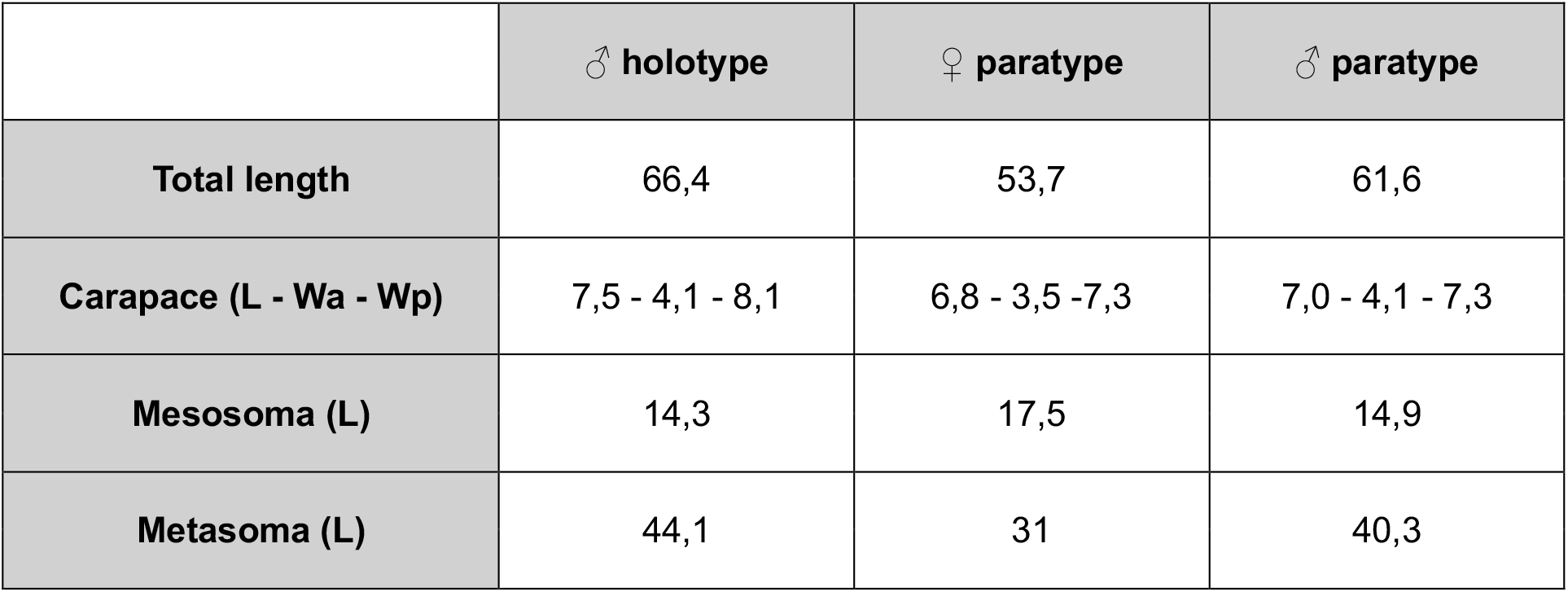
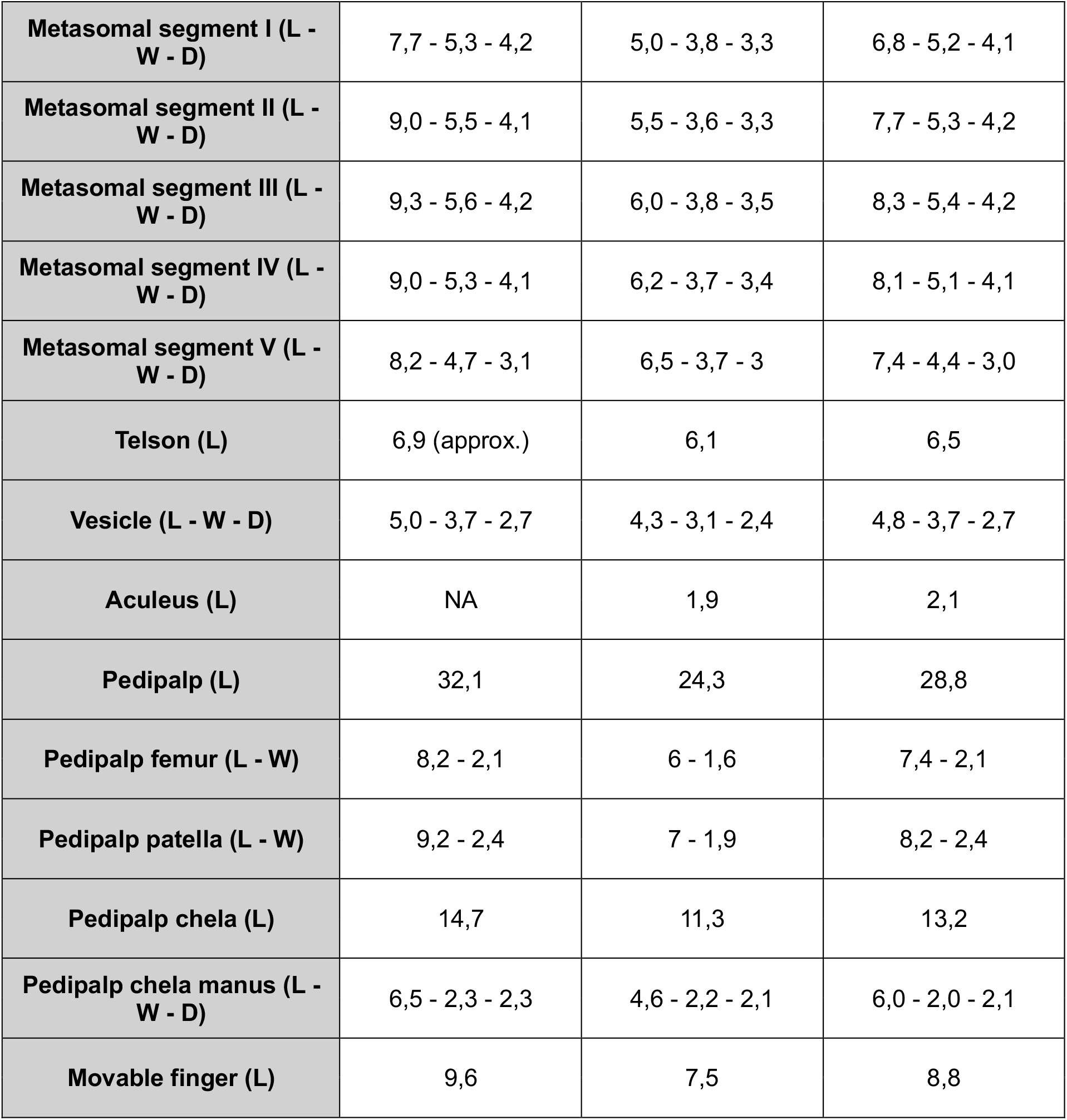
Morphometric measurements for the type specimens of *T. (Tityus) XXX* sp. nov. Measurements are given in mm. The aculeus of the male holotype is broken and could not be measured. Abbreviations: L (Length); W (Width); D (Depth); a (anterior); p (posterior).

|  | ♂ holotype | ♀ paratype | ♂ paratype |
| --- | --- | --- | --- |
| **Total length** | 66,4 | 53,7 | 61,6 |
| **Carapace (L - Wa - Wp)** | 7,5 - 4,1 - 8,1 | 6,8 - 3,5 - 7,3 | 7,0 - 4,1 - 7,3 |
| **Mesosoma (L)** | 14,3 | 17,5 | 14,9 |
| **Metasoma (L)** | 44,1 | 31 | 40,3 |
| **Metasomal segment I (L - W - D)** | 7,7 - 5,3 - 4,2 | 5,0 - 3,8 - 3,3 | 6,8 - 5,2 - 4,1 |
| **Metasomal segment II (L - W - D)** | 9,0 - 5,5 - 4,1 | 5,5 - 3,6 - 3,3 | 7,7 - 5,3 - 4,2 |
| **Metasomal segment III (L - W - D)** | 9,3 - 5,6 - 4,2 | 6,0 - 3,8 - 3,5 | 8,3 - 5,4 - 4,2 |
| **Metasomal segment IV (L - W - D)** | 9,0 - 5,3 - 4,1 | 6,2 - 3,7 - 3,4 | 8,1 - 5,1 - 4,1 |
| **Metasomal segment V (L - W - D)** | 8,2 - 4,7 - 3,1 | 6,5 - 3,7 - 3 | 7,4 - 4,4 - 3,0 |
| **Telson (L)** | 6,9 (approx.) | 6,1 | 6,5 |
| **Vesicle (L - W - D)** | 5,0 - 3,7 - 2,7 | 4,3 - 3,1 - 2,4 | 4,8 - 3,7 - 2,7 |
| **Aculeus (L)** | NA | 1,9 | 2,1 |
| **Pedipalp (L)** | 32,1 | 24,3 | 28,8 |
| **Pedipalp femur (L - W)** | 8,2 - 2,1 | 6 - 1,6 | 7,4 - 2,1 |
| **Pedipalp patella (L - W)** | 9,2 - 2,4 | 7 - 1,9 | 8,2 - 2,4 |
| **Pedipalp chela (L)** | 14,7 | 11,3 | 13,2 |
| **Pedipalp chela manus (L - W - D)** | 6,5 - 2,3 - 2,3 | 4,6 - 2,2 - 2,1 | 6,0 - 2,0 - 2,1 |
| **Movable finger (L)** | 9,6 | 7,5 | 8,8 |

### -T. charalaensis

i. coloration of the metasoma (yellowish in *T. charalaensis* and reddish in *T. XXX* sp. nov.)
ii. lateroventral carinae of metasomal segment II occupying its distal two-thirds (distal fifth in *T. XXX* sp. nov.)
iii. lower pectinal tooth count with 14 in female holotype (15-16 in *T. XXX* sp. nov.)
iv. 14 rows of denticles on movable chela finger (15 in *T. XXX* sp. nov.)
v. female pedipalp femur narrower with length/width ratio 4,71 (3,75 in *T. XXX* sp. nov.)
vi. female pedipalp patella broader with length/width ratio 3,09 (3,68 in *T. XXX* sp. nov.)
vii. other morphometric values, see **table 2**

### Ecology and type locality

The type locality, located in Cundinamarca, Colombia, is part of the Magdalena valley montane forest ecoregion. All specimens were collected on a patch of medium-altitude broadleaf rainforest, 1350 m a.s.l., with a mean annual precipitation of approx. 1450mm and mean annual temperature of 21.3°C. The new species occurs in sympatry with *T. (Atreus) icarus* (Laborieux, 2024), but with clear differences in microhabitat preferences. While the latter species is strictly arboreal, usually found hanging vertically on tree trunks, specimens of *T. XXX* sp. nov. were collected on the forest floor exclusively, with no observed arboreal tendencies. Adult specimens were exceedingly rare, perhaps suggesting ontological variation in microhabitat preferences.

## 4 Discussion

The new taxon described herein represents the latest addition to Colombia’s already remarkable scorpion diversity. *Tityus XXX* sp. nov. is the 229th species described in genus *Tityus*, and its discovery in a relatively well-prospected region (Cordillera Oriental) highlights the importance of comprehensive geographic sampling for taxonomic research. While *Tityus* species are responsible for a considerable proportion of the scorpionism cases reported from Colombia (Chippaux & Goyffon, 2008; Otero et al., 2004), the epidemiological implications of a new venom-spraying species are unclear. On the one hand, *T. XXX* sp. nov. occurs in an area with relatively high population density for the country (approx. 90 inhabitants/km^2^ according to DANE, 2005), increasing the risk of contact between scorpions and human individuals. On the other, the range of venom sprays is moderate, with a maximum estimate of approximately 36 cm. It is thus likely that such an attack would only reach human eyes in a situation where a direct sting (*e*.*g*., to the hands) is also a threat. There does not appear to be any record of visual impairments following scorpion-related accidents in the country, providing no evidence for the involvement of venom spraying in Colombian scorpionism. Overall, the risk presented by *T. XXX* sp. nov. may be restricted to the venomous stings it can deliver.

The contrasting ballistic characteristics of venom flicks and venom sprays point to distinct mechanisms of projection. The venom sprays of *T. XXX* sp. nov. are likely analogous to those of *P. transvaalicus*, where rapid contraction of venom gland muscles has been speculated to propel the secretion at high speed (Newlands, 1974; Nisani & Hayes, 2015). Unfortunately, the exact expulsion mechanism of *Parabuthus* venom pulses has not yet been investigated, and how much functional specialization is required to achieve venom spraying is unknown. Perhaps more enigmatic are the venom flicks, whose considerably lower volumes and velocity strongly suggest a sting-like mechanism. Specifically, the mean volume estimated for flicks is consistent with values previously reported for stings from similarly-sized scorpions (*Smeringurus mesaensis* Stahnke, see Van Der Meijden et al. (2015)), and close inspection of the high-speed videos suggests that much of the velocity of flicked venom is contributed by metasomal movements. If flicks represent sting-like volumes of venom, and if the contraction of the venom gland does not contribute significantly to the velocity of the toxungen, venom flicking may be mechanically equivalent to a mid-air sting. One key difference between flicks and stings, however, is that venom flow rate measurements made by Van Der Meijden et al. (2015) are greatly exceeded by expected flick flow rates. While stung and flicked volumes may be comparable, reported injection times during stings were 34 – 95x longer than the expulsion time considered here for venom flicks. Elucidating the biomechanical underpinnings of venom pulses in *T. XXX* sp. nov. may thus require the characterization of its own stinging behaviour, and improved understanding of the muscles responsible for venom gland contraction. Another important consideration would be the acquisition of direct volume measurements for venom pulses. Although mathematically sound, the volumes presented here are secondary estimates (*i*.*e*. depending on estimated stream width, pulse velocity and duration) and should only be treated as exploratory.

In addition to their distinct ejection velocities, venom flicks and sprays may be functionally specialized through differences in the initial position and orientation of the telson. Trends in the data suggest that, on average, venom flicks are initiated downward and from a lower position than venom sprays, which tend to be initiated at a positive angle (*θ*_*i*_> 0°). While certainly commanding caution, the weak statistical support for these results could stem from relatively small sample sizes and high variance. If true, the observed patterns of angular and vertical positioning reaffirm the role of sprays as an optimized ranged attack, but inferring the biological function of venom flicks remains challenging. The estimated range and vertical reach of venom sprays are consistent with known *Tityus* predators such as amphibians (Botero-Trujillo, 2006; Jared et al., 2020), birds (Murayama et al., 2022) and lizards (Silva-Júnior et al., 2023). Larger animals, otherwise out of reach, may also expose themselves by lowering their heads to the ground while attempting predation. Conversely, a hypothesis based on the range and downward trajectory of flicks could be that they target not the eyes, but the respiratory tissues of a potential predator. When directly facing a scorpion, the snout or beak of an antagonist would likely be significantly closer and lower than its eyes, increasing its potential exposure to venom flicks. The low volume of the flicked droplets would, in that case, provide a cheap alternative to spraying while facilitating inhalation. Human respiratory irritation has been reported following *P. transvaalicus* venom sprays (Nisani & Hayes, 2015), lending credibility to this interpretation. Alternatively, venom flicking in *T. XXX* sp. nov. may be triggered by an uncontrolled nervous impulse and contraction of the venom gland under stress, serving no true biological purpose. The metabolic cost of venom (Evans et al., 2019; Morgenstern & King, 2013) and the apparent absence of such a phenomenon in *Parabuthus* make this an unlikely possibility.

Overall, three key differences with *P. transvaalicus* suggest that the venom spraying behaviours of both species evolved to face different selection pressures. First, *T. XXX* sp. nov. employs venom pulses in response to stimuli of lower intensity than those triggering the behaviour in *P. transvaalicus*. Important work by Nisani & Hayes (2015) has shown that the latter species did not spray venom when prodded on the legs or body, only doing so when their metasoma was grasped. Interestingly, the incidence of venom spraying was then dictated by perceived threat levels, remaining low (adults) to null (juveniles) when grasping was the only stimulus and increasing significantly with the addition of airborne (pressurized air) stimulation. In juvenile *T. XXX* sp. nov., however, applying light pressure to the carapace was enough to obtain high venom pulse rates. Further, the metabolic cost of venom pulses is likely dramatically different between both species. *P. transvaalicus* produces low quantities of prevenom, representing approx. 5% of the total by volume (Inceoglu et al., 2003). This small reserve may be exhausted in a single sting, and venom sprays invariably use metabolically costly venom. The new species, in contrast, possesses a comparatively large reserve of prevenom-like secretion, which it can use in repeated stings and venom pulses as the threat persists. *T. XXX* sp. nov. still produces opaque venom, similar to *P. transvaalicus* (Inceoglu et al., 2003) and other *Tityus* species (Lira et al., 2017), but its use appears to be unusually rare. In a study on *Tityus stigmurus* Thorell, individuals expelled white venom 93% of the time in a situation representing an equivalent, if not lower, threat level to that presented to the *T. XXX* sp. nov. trial group (Lira et al., 2017). The term “prevenom” may therefore be misleading in the case of the new species, since this clear mixture could in fact support both defense and predation as a primary secretion. Finally, both spitting species differ in their initial spray velocities, with that of *T. XXX* sp. nov. being almost twice as high on average as that of *P. transvaalicus*. (Nisani & Hayes, 2015). Overall, these differences support an interpretation depicting *Tityus* venom sprays as a highly specialized and relatively cheap defensive strategy used in frequent antagonistic encounters, with an emphasis on range, while those of *Parabuthus* may represent a last resort against a large predator having already grasped the scorpion’s metasoma.

In conclusion, the discovery of a new venom-spraying scorpion in South America constitutes an exciting new case of convergent evolution, calling for further exploration. For example, this study only considered juveniles due to the unavailability of mature specimens, but adults are expected to employ similar defensive strategies. Comparing venom spraying across developmental stages may provide new insights into the ecological pressures driving the evolution of such an uncommon behaviour. Future studies should aim to produce precise quantitative data on venom expenditure through the direct collection of projected secretions, while investigating their toxicology. At a broader scale, a direct comparison of *Parabuthus, Tityus* and *Hadrurus* venom spraying may expose interesting patterns of behavioural or chemical adaptation, advancing our understanding of venom modulation, specialization, and evolution.

## 5 Acknowledgements

I would like to thank Carolina Pardo Díaz (Universidad del Rosario, Colombia) for her continued support in handling administrative processes and arranging permits. I am also indebted to Daniela Lozano-Urrego and Geraldine Rueda for their support and affectionate company during my stay in Colombia. In addition, I am deeply grateful to Gonzalo Giribet and Eric Ythier for their guidance in preparing this manuscript and a number of stimulating discussions that helped improve it. Finally, I would like to express my sincere appreciation to Arie van der Meijden, Tate Yawitz and Rowan Stanforth for their helpful comments on theoretical and experimental matters. All specimens were legally collected under a permit issued to Universidad del Rosario by ANLA (*Permiso Marco de Recolección de Especímenes de Especies Silvestres de la Diversidad Biológica con Fines de Investigación Científica No Comercial*, #0530).

## Notes

### Competing Interest Statement

The authors have declared no competing interest.

